# Comparison of rule- and ordinary differential equation-based dynamic model of DARPP-32 signalling network

**DOI:** 10.1101/2022.03.26.485918

**Authors:** Emilia M. Wysocka, Matthew Page, James Snowden, T. Ian Simpson

## Abstract

Dynamic modelling has considerably improved our understanding of complex molecular mechanisms. Ordinary differential equations (ODEs) are the most detailed and popular approach to modelling the dynamics of molecular systems. However, their application in signalling networks, characterised by multi-state molecular complexes, can be prohibitive. Contemporary modelling methods, such as rule-based (RB) modelling, have addressed these issues. The advantages of RB modelling over ODEs have been presented and discussed in numerous reviews. In this study, we conduct a direct comparison of the time courses of a molecular system founded on the same reaction network but encoded in the two frameworks. To make such a comparison, a set of reactions that underlie an ODE model was manually encoded in the Kappa language, one of the RB implementations. A comparison of the models was performed at the level of model specification and results were acquired through model simulations. Conforming to previous reports, we confirm that the Kappa model recapitulated the general dynamics of its ODE counterpart with minor differences. These differences occur whenever molecules have multiple sites binding the same interactor. Furthermore, activation of these molecules in the RB model is slower than in the ODE one but can be corrected by revision of the rate constants used in the relevant rules. As in previous reports on other molecular systems, we find that, also in the case of the DARPP-32 reaction network, the RB representation offers a more expressive and flexible syntax that facilitates access to fine details of the model, facilitating model reuse. In parallel with these analyses, this manuscript reports a refactored model of the DARPP-32 interaction network that can serve as a canvas for the development of a more complex interaction network to study this important molecular system.

## INTRODUCTION

Computational dynamic modelling frameworks probe mechanistic and quantitative aspects of molecular interactions, which can grant the development of mechanism-based therapies with more predictive power on outcomes of therapeutic interventions (Xie et al., 2014). Such interventions often target molecular signalling (Li and Mansmann, 2014; Jia et al., 2019; Volkow and Boyle, 2018), characterised as a complex system of coupled interacting components, often proteins, forming networks that activity leads to non-additive effects (Kitano, 2002). Defining molecular reactions as a set of coupled ODEs has traditionally enabled dynamic modelling of molecular pathways (Sible and Tyson, 2007). ODE-based modelling is a powerful and acclaimed formalism with a long tradition. It has established standards, and various software tools that support and facilitate the formulation and analysis of ODE models (Dräger et al., 2008; Hoops et al., 2006; Sible and Tyson, 2007). However, the explicit enumeration of molecular species as variables required by the equation-based formulation excludes the representation of molecules that assemble into multivalent protein complexes; typically these have multiple functionally divergent states, a common characteristic of molecules involved in cell signalling (Hlavacek et al., 2003). Currently developed computational modelling methods effectively addressed these challenges of expressivity and increasing complexity of the modelled systems. An example of such a method is RB modelling, designed to model interacting proteins. Compared to the ODE-based paradigm, which represents the molecular system as concentrations of molecular species and focuses on their reaction kinetics, RB modelling is an agent-centred method in which the distribution of molecular compositions can be studied along with their abundance (Faeder et al., 2003; Blinov et al., 2004; Danos and Laneve, 2004). The potential of the RB paradigm has been extensively discussed (Danos, 2007; Chylek et al., 2013, 2014, 2015) and demonstrated with examples (Danos, 2007) or with models answering novel biological questions across signalling, regulatory and metabolic networks (Wilson-Kanamori et al. 2015; Antunes et al. 2016; Di Camillo et al. 2016; Santibáñez et al. 2020; Chattaraj et al. 2021; Nosbisch et al. 2022; see reviews for earlier models published before 2006 - Hlavacek et al. 2006; and from 2007 to 2013 - Chylek et al. 2014). These models are often based on chemical reaction networks previously developed for ODEs. Though the RB paradigm was implemented to match the concept of chemical reactions, rules are generators of reaction networks that can potentially lead to diverging results. Thus, as the modelling paradigm can affect the underlying model specification, it would be informative to compare simulations of RB and ODE models defined by the same reaction network, originally built to be solved with ODEs. RB and ODE models have been compared on various aspects before. For instance, Blinov et al. (2006) compared network-like model of epidermal growth factor receptor (EGFR) based on reactions first defined in the ODE model of Kholodenko et al. (1999), with simplified pathway-like structure. However, several assumptions underlying the original model were purposefully modified what changed the underlying interaction network, such as the dissociation of EGFR dimers when phosphorylated or bound to other molecules. In addition to contrasting rules to reactions, another intensively studied aspect, finalised with positive conclusions, was whether stochastic simulation can reveal new properties of a system previously modelled as deterministic one (Vlysidis and Kaznessis, 2018; Hahl and Kremling, 2016; Bustos et al., 2018). As simplistic as it may sound, none of the comparisons mentioned were intended to painstakingly disassemble reaction using the rules of a relatively medium-sized model and compare simulation results. Thus, this study compares the existing ODE model with the new RB model, where both models are based on the same definition of chemical reactions. We specifically ask: (1) how closely the dynamics of an ODE-based model can be replicated with an RB model? (2) if differences between the two are observed, what is the underlying cause? (3) if the dynamics are indeed replicated, what advantages are there to using an RB model? We present the results of the models’ comparison at the level of notation and using model dynamics under different conditions. The advantages and disadvantages of the two model representations are discussed, alongside suggestions for future research. We hope that this small-scale attempt may nevertheless bring value to the modelling community to better choose between the two different approaches.

## BACKGROUND

Dynamic computational modelling frameworks consist of model specification schema and simulation methods. The model specification is a set of equations or instructions written in a machine-interpretable language. These languages are based on mathematical formalisms that define relationships between variables whose quantities vary over time. Models are run as numerical simulations using algorithms that interpret the model’s mathematical formulation and calculate changes in the quantities of model variables. By adopting a suitable level of abstraction and with the use of a sufficiently expressive language, systems can be modelled such that experimentally-derived evidence can be incorporated to improve the model’s quality. A formal approach to the generation of models is desirable to (i) encode facts in an unambiguous and explicit manner, (ii) facilitate the understanding of models, (iii) allow easy modification of models to accommodate more than one hypothesis, (iv) aid interpretation of the underlying biological phenomena, and (v) provide a standard approach to the integration of novel data sources (Kitano, 2002). These features are especially important because model generation often requires knowledge spanning multiple disciplines; the existence of formal modelling frameworks enforces a common understanding of the explicit meaning of model components.

In the first step of building a kinetic model, transitions from reactants into products are defined as *chemical reactions* between molecules. Quantitative evaluation of model behaviour over time has been commonly achieved by converting coupled chemical reactions to a set of ODEs that are solved with numerical procedures (Wilkinson, 2006). Each rate equation expresses the change of concentration of a single molecular species over time, formulated with reaction rates that directly take part in the creation and elimination of this species (Sible and Tyson, 2007; Klipp et al., 2005; Hlavacek et al., 2006). Each reaction rate is weighted by a reaction-specific *rate constant*. Time courses obtained by solving ODE models are continuous and deterministic, characterised by smooth and gradual change of species concentrations over time (Wilkinson, 2009). Although, this setup does not reflect the actual characteristics of subcellular events driven by random collisions between discrete molecules (Gillespie, 1976), this approach is correct as long as abundances of reactants are large enough to render random fluctuations as negligible (Chen et al., 2010). If this condition is violated, which may be the case in synaptic signaling networks, the current development of machine-readable formats for encoding biological models (e.g. Systems Biology Markup Language (SBML)) makes it possible to obtain trajectories of the same model using deterministic solvers and stochastic simulators (Gillespie, 1977; Hoops et al., 2006). A more critical shortcoming of ODE-based models lies in the requirement for explicit enumeration of all molecular species in signalling networks (Hlavacek et al., 2003; Danos, 2007). This drawback precludes mechanistic modelling of systems with multi-state promiscuous molecules that can adopt combinatorially complex states (Seshacharyulu et al., 2012; Chen et al., 2016; Mayer et al., 2009; Suderman and Deeds, 2013), where only a small fraction can be represented with ODEs (Chylek et al., 2014). However, the development of formal methods in computer science has expanded the number of observed properties of biological systems that can be dynamically and quantitatively modelled (Bartocci and Lió, 2016) (reviewed in Machado et al. 2011; Tenazinha and Vinga 2011; Bartocci and Lió 2016; Ji et al. 2017; Le Novère 2015). As this study examines an alternative to ODEs, the focus lies on non-spatial single-scale mechanistic methods. Of those that fit this characterisation (Baeten, 2005; Ciocchetta and Hillston, 2008; Regev et al., 2001; Guerriero et al., 2007; Dematté et al., 2010) we chose rule-based modelling (RBM) as a suitable framework for representing systems with multistate combinatorial interactions. RBM represents molecules are structured graph objects (nodes with sites) and reactions as graph transformations, where bonds are formed between the node sites. Graph transformations are encoded as rules that are instructions for local and conditioned transformations. Therefore, a rule represents either a set of reactions or an exact reaction instance, dependent on details of said conditions. By this, a rule can express an infinite number of reactions with a small and finite number of generalised rules (Chylek et al., 2013). A protein abstracted as a node with binding sites that have internal states gives a sufficiently expressive system to capture the principal mechanisms of signalling processes (e.g. biding, dissociation, synthesis, degradation, and state change; Liu and Thiagarajan 2012), providing insight into site-specific details of molecular interactions (e.g. affinities, dynamics of post-translational modifications, domain availability, competitive binding, causality) and the intrinsic structure of interaction networks. Among two major RBM implementations (Stefan et al., 2014), the very first being BioNetGen (BNG) (Faeder et al., 2003; Blinov et al., 2004; Faeder et al., 2009), the Kappa framework (Danos and Laneve, 2004) was chosen for this comparison. Although there are some notational differences between the two frameworks, the reactions are coded as rules in virtually the same way (Suderman and Hlavacek, 2017). In addition, Suderman and Deeds (2013) reported that the two frameworks produced exactly the same dynamics for compared models. The language consists agents, patterns, rules and observables (Feret et al., 2009a). An *agent* represents a reactant with *interface*, that has a set of *sites* that express *internal state* and *binding state* (Feret et al., 2009a). Agents during the system simulation form *mixture* of *molecular species* that contain a full description of their states and site occupancy. Agents can also have a partially specified interface, i.e. expressed as a *pattern* (Feret et al., 2009b), thereby matching larger groups of molecular species. *Rules* define state transformations or actions of agents, i.e. reactions. Dependent on the generality of agent description, a single rule can represent a set of reactions. On the other hand, a rule specified with complete information about the agent’s interface is in one-to-one correspondence with a reaction. *Observables* are declared variables whose time courses are the result of a simulation. The trajectory of an observable declared as an incomplete pattern is the sum of the trajectories of the many molecular species that fit the pattern. The Kappa language also provides means to induce perturbations during the simulation. Perturbations can be applied once or repeatedly according to the indicated conditions, to update rate constants, or add molecules. *Snapshots* of a molecular mixture, taken during the simulation, provide information on molecular species and their quantities at specified time-points. KaSim is the Kappa simulator (Danos, 2007; Krivine et al., 2009), based on Gillespie’s Stochastic Simulation Algorithm (SSA) (Gillespie, 1976, 1977) that numerically simulates individual stochastic trajectories of molecules defined with reaction networks (Gillespie, 1977; Wilkinson, 2009) whose efficiency is independent of the network size (Yang and Hlavacek, 2011).

To directly compare models’ simulations, we selected an ODE model available in a machine-readable format that can be numerically simulated in non-obsolete software, thus satisfying the reproducibility criterion. A model of the immediate interactors of dopamine- and cAMP-regulated neuronal phosphoprotein with molecular weight 32 kDa (DARPP-32) network by Fernandez et al. (2006) satisfies this requirement. The model is also considered a valuable study that can serve as a solid core for the construction of larger and more complex models in the community interested in the modelling of dopamine-dependent synaptic plasticity (Manninen et al., 2011). Finally, it is a study widely cited not only by modellers (Nair et al., 2016; Nakano et al., 2010; Mattioni and Le Novère, 2013) but also by experimentalists (Bales et al., 2011; Kim et al., 2015; Buesa et al., 2016). It should be noted that the stochastic simulation for this particular system should not show any significant differences from the ODE model, as the number of particle copies is sufficiently large.

### DARPP-32 and its interaction network

DARPP-32, officially named as Protein Phosphatase 1 Regulatory Inhibitor subunit 1B isoform 1 (PPP1R1B) (NCBI Resource Coordinators, 2016), is an important multistate and intrinsically disordered protein (Marsh et al., 2010) regulating dopamine-dependent synaptic plasticity in medium spiny projection neurons (MSPN) of the striatum. Around 95% of human striatal cells are MSPNs, in which signalling cascades activated simultaneously by glutamatergic and dopaminergic stimuli are a necessary condition for the long-term potentiation (LTP) that underlies context and reward-related learning (Beninger and Gerdjikov, 2005). Furthermore, DARPP-32 malfunction and abundance relate to multiple central nervous systems (CNS) disorders. Among these are Alzheimer’s disease (Cho et al., 2015), addiction (Philibin et al., 2011), affective disorders (Kunii et al., 2014), and schizophrenia (Kunii et al., 2014; Wang et al., 2015). Glutamatergic and dopaminergic signals are integrated by DARPP-32 which is involved in a complex network of interactions regulating its multiple phosphorylation sites, of which 4 are known to have a regulatory impact on DARPP-32 itself (Yger and Girault, 2011): Threonine 34 (Thr34), Threonine 75 (Thr75), Serine 137 (Ser137), and Serine 102 (Ser102). The multiplicity of phosphorylation states leads to numerous interacting partners for DARPP-32 that affect these states. The Threonine sites (Thr34, Thr75) have major regulatory roles in signal processing. The Serine sites (Ser137, Ser102) have a supporting role in Thr34 signal enhancement.

Fernandez et al. (2006) studied the integrative effect and sensitivities of dopamine (DA)- and glutamate (Glu)-mediated signals on the DARPP-32 network (Figure 1). Their model examined the particular effect of cyclic adenosine monophosphate (cAMP)-pulse followed by calcium ions (Ca^2+^) spike trains whilst varying the distance between the stimuli. The study showed that DARPP-32 is a robust integrator, indifferent to its initial concentration and delay between the stimuli, far more complex than a bistable switch between DA and Glu signals. To reproduce the system’s behaviour, the authors included two main pathways that mediate these signals, cAMP–PKAxydDARPP-32 phosphorylated at Threonine 34 (D34) and Ca^2+^–PP2B–DARPP-32 phosphorylated at Threonine 75 (D75). Contrary to the majority of previous models of Glu and DA signal integration (Lindskog et al., 2006; Gutierrez-Arenas et al., 2014; Nair et al., 2014, 2016), DARPP-32 included three phosphorylation sites: Thr34, Thr75 and Ser137. The authors performed two *in silico* mutagenesis experiments modifying the role of Ser137. The first mutation inhibits site phosphorylation by changing Serine to Alanine at the 137 position (Ser137Ala). The second mutation leads to permanent phosphorylation of the Ser137 site (constSer137).

**Figure 1.**
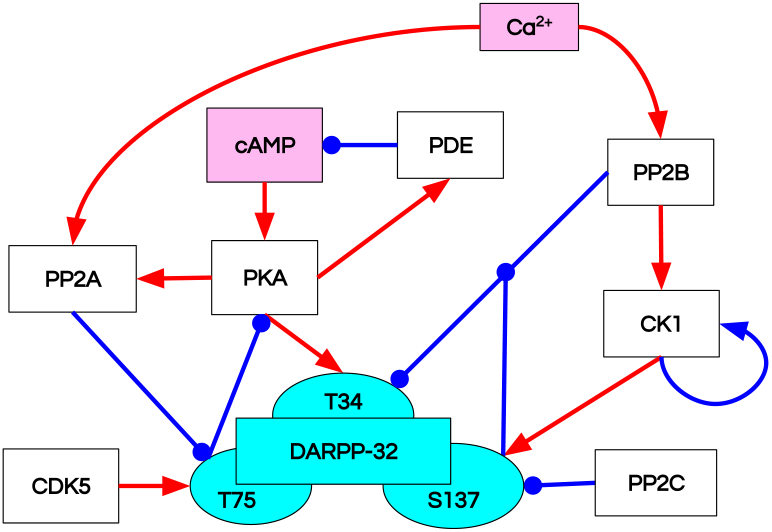
Reaction diagrams representing different aspects of the DARPP-32 network included in the ODE model by Fernandez et al. (2006) Nodes: DARPP-32 (*cyan*), second messengers (*magenta*), kinases/phosphatases (*white*). Edges: inhibition reactions (*blue*), activation reactions (*red*).

## METHODS

The first step of comparison of RB and ODE frameworks involved encoding reactions underlying the ODE model of Fernandez et al. (2006) (“model B”) into the Kappa language, version 3.5 (Feret and Krivine, 2012). Then, models were simulated in different variants to obtain time courses of equivalent observables to compare (Figure 2).

**Figure 2.**
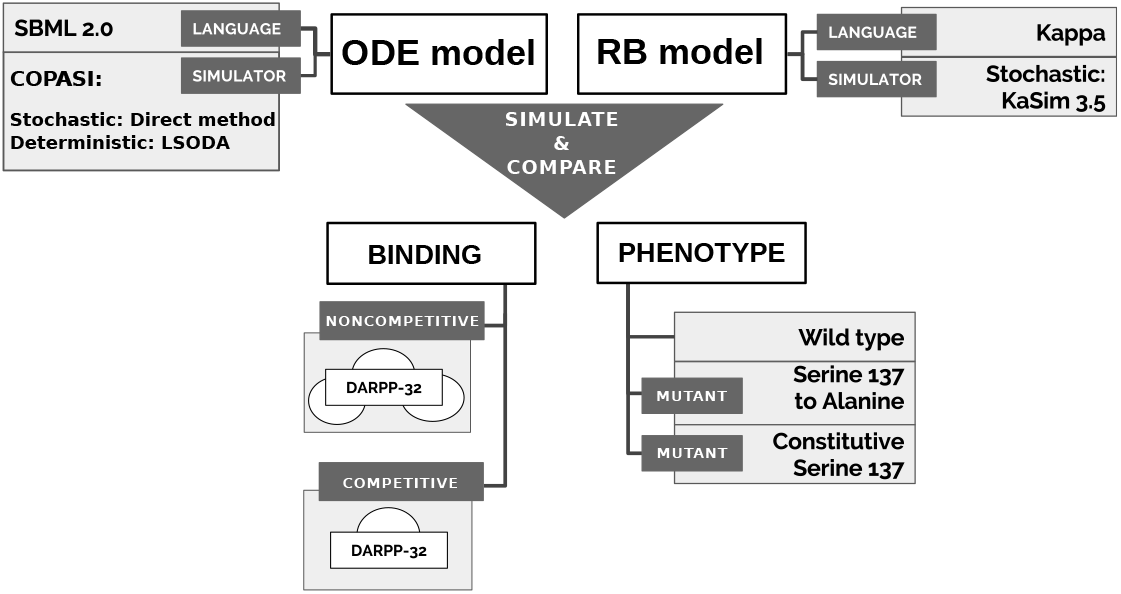
Approach to comparison of ODE and RB modelling frameworks.

### Model translation

As a single reaction can be written as a single rule, the translation could have been accomplished in a one-to-one manner. However, to fully benefit from rule patterns, multiple reactions can be condensed into fewer rules by removing irrelevant context, i.e. *decontextualised*. The context of a reaction in the RB model is defined as the information about the agent’s binding sites, partners and internal states. Based on this definition of reaction context, the following criteria guided decisions about condensing reactions into rules. Given a set of reactions of the same type (forward, backward, or catalytic) between the same reactants (agents), if the difference between reactions lies in agent states (internal or binding) that do not change after the transition from reactants to products, and reaction constants (rates) in all these reactions have the same values, then information about agent states does not define reaction conditions; hence, it can be removed from the reaction notation, i.e. a set of reactions become a rule pattern.

Among the least intuitive cases in encoding reactions into rules are complex substrate activations of PKA and PP2B. PKA is activated by the binding of 4 cAMP molecules, whereas PP2B activation requires 4 Ca^2+^ ions. In other words, multiple molecules of the same type bind substrates on different sites that have to be uniquely named, which requires explicit encoding of all possible binding combinations on 4 different sides (approach after Danos et al. (2011)), called hereafter as *combinatorial binding*.

We also need to translate molecular concentrations, rate constants and initial molecular abundances to copy-numbers (Feret and Krivine, 2012; Sekar and Faeder, 2012). Lastly, the cAMP pulse and the Ca^2+^ spiking are reproduced by the addition of molecular copy numbers and modification of rate constants during the simulation, respectively.

### Approach to comparison of models

The qualitative comparison of the results of two dynamic models required the simulation of both models in a stochastic scheme and the alignment of the trajectory of the corresponding observables under varying conditions to allow more comprehensive comparisons between the frameworks (Figure 2).

#### Selecting and pairing observables

The plots in the original publication show aggregated variables that are summed trajectories of multiple molecular species. For instance, “D34” denotes DARPP-32 phosphorylated at Thr34, regardless of its state of binding or other phosphorylation sites. The concept of aggregated variables corresponds to *observables* in RB modelling, and therefore, we use the term *observable* hereafter to denote aggregated variables. Observables of the ODE model were aggregated here based on their names matching partial strings representing the observables of interest. To verify this approach, obtained observables of the ODE model with this method were qualitatively compared with the 6 observables plotted in the original publication. The choice of other observables follows these principles: (1) if an agent has internal states, the activated state is set as its observable form, e.g. “CK1u”; (2) if an agent is created and degraded over the simulation, its observable is set to its least specific form, e.g. “PKA”; (3) if an agent is not created and degraded during the simulation, i.e. its level remains constant throughout the simulation and has no internal states, its observable is set to its bound form, e.g. “_CDK5” (see Table 1 for the complete list of RB and ODE observables with definitions).

**Table 1.** The specification of a model can be divided into components. The total number of elements in each component is shown for both the ODE model and the RB model.

| ODE model |  |  | RB model |
| --- | --- | --- | --- |
| Model Component | Total counts | Total counts | Model Component |
| Reaction instances | 152 | 132 | Reaction rules |
| Concentration-based rate constants | 152 | 62 | Stochastic rate constants |
| Initial concentrations | 75 | 8 | Initial copy numbers |
| Molecular species | 75 | 91/137 | Molecular species |
| Stimuli events | 21 | 21 | Stimuli events |

#### Model simulation

We simulate both models in the stochastic scheme but within different simulation environments. The RB model was simulated with KaSim. The SBML format of the ODE model was run with COPASI (version 4.20), a common simulation environment for SBML-formated models (Hoops et al., 2006), using the deterministic solver (LSODA) and stochastic simulator (implementation of *direct method* by Gillespie (1977)).

#### Model perturbations

The first type of perturbation is based on a modification of rate constants. We used this modification to induce site-directed mutations: Ser137Ala and constSer137, replicating the original study (Fernandez et al., 2006). In both cases, the alteration of the model involved the inactivation of 4 reactions by zeroing their rate constants. In the RB model, they are represented by 1 rule, i.e. a change of a single constant induced each mutagenesis.

We additionally tested the RB model with two different binding schemes, applicable only to the RB model, and further called noncompetitive and competitive binding (Figure 2). In the noncompetitive binding, all interactors of DARPP-32 can bind simultaneously to three different sites. The competitive binding assumes one interaction with DARPP-32 at a time, which reflects the ODE model assumption. The Fernandez et al. (2006) study does not discuss if DARPP-32 binding partners bind to the same or different active sites, though DARPP-32 is an intrinsically disordered protein with an unknown binding interface (Dancheck et al., 2008; Mollica et al., 2016; Engmann et al., 2015; Choy et al., 2012). The ODE model specification demonstrates that DARPP-32 forms at most heterodimers. This type of modification in the ODE model would require enumeration of additional molecular species, the addition of new equations and updating the existing ones. Contrary to the ODE model specification, a definition of such a binding scenario in the RB notation requires the same number of rules, provided that concurrently bound interactors do not influence each other.

## RESULTS

Comparison of models was performed on two levels, model notation and simulation results. The model notation was analysed by dividing the model into components and comparing their sizes. We expect the set of reactions underlying the ODE model to be represented with fewer rules since a single one can constitute a pattern representing several reactions. The comparison of simulation results involved the alignment of equivalent time courses obtained by model simulations. We performed the comparison of time courses between three variants of each model: (1) base-line condition (wild-type) and two site-directed mutations: (2) Ser137Ala and (3) constSer137. Finally, we compared two RB model variants, representing: (1) DARPP-32 with a single binding site; and (2) DARPP-32 with three independent binding sites.

### Rule patterns reduce reaction number in a certain type of model components

Table 1 juxtaposes the total counts of model components in each model. The RB model represents 152 reactions with 132 rules, each parameterised by one of 62 unique rate constants. This number is lower than the total number of rate constants used to parameterise the ODE model (152). The final rule set is more than twice as large as the unique number of rate constants, meaning that more than one rule is parameterised by the same rate constant. The number of molecular species in the RB model, obtained with snapshots capturing the state of the molecular mixture over simulation time (every 10000th event), is 91 for the competitive RB model, and 137 for the non-competitive one. In both cases, the sum of molecular species is higher than in the ODE model (75). As expected, the number of rules corresponding to reactions is lower, and the number of molecular species is much higher, confirming that expression patterns reduce the number of rules needed to represent a reaction system. Nonetheless, the number of rules is only slightly lower than the number of reactions (152 to 131). If we closely compare models by parts representing more general molecular mechanisms, rule representation reduces the reaction number in some components but extends it in others (Table 2). The reduction occurred only in “DARPP-32 phosphorylation” and “PP2A activation by Ca^2+^” components, where combinations of states of DARPP-32 phosphorylation sites do not have to be explicitly named. In contrast, the increase in the number of reactions compared to the reactions occurred in the components “PKA activation” and “PP2B activation”. They both have 4 sites that bind the same molecules, Ca^2+^ and cAMP, respectively, which requires the expression of combinatorial binding in the rule notation.

**Table 2.**
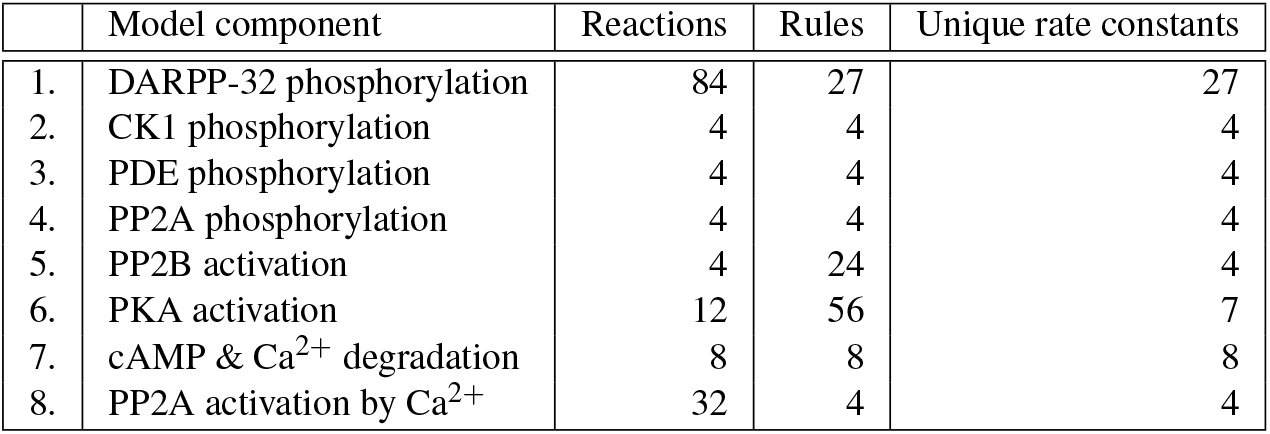
The list of reactions in the Fernandez et al. (2006) publication was divided into components based on more general molecular processes represented by subsets of reactions, such as phosphorylation or activation. We can closely examine reaction-rule relation by comparing models by components. The table shows the number of reaction rules versus reaction instances and a unique number of rates per model component. It is noticeable that the reduction in the number of reaction instances due to the translation of reactions into Kappa language occurred in only two model components (1. & 8.), while in two others, it resulted in an expansion of the rule number (5. & 6.).

### RB model recapitulates dynamics of ODE model with minor discrepancies

The RB model recapitulates the principal dynamics of the ODE model, albeit there are some observable differences (compare Figures 3B and C). For instance, there is a comparable variability of trajectories of Thr34 phosphorylated isoform (“D34” and “D34*”) that can be observed in the relaxation phase (after the 600th time point). “D34” in the ODE model needs 100 more time steps to reach the second peak, and it is weaker than its RB counterpart (“D34*”). Worth noting is that the standard deviation in the stochastically simulated ODE model reveals a distinctive variation in abundance of the “D34” observable during the relaxation phase. We further use the stochastic trajectories of the ODE model for comparison with the RB model.

**Figure 3.**
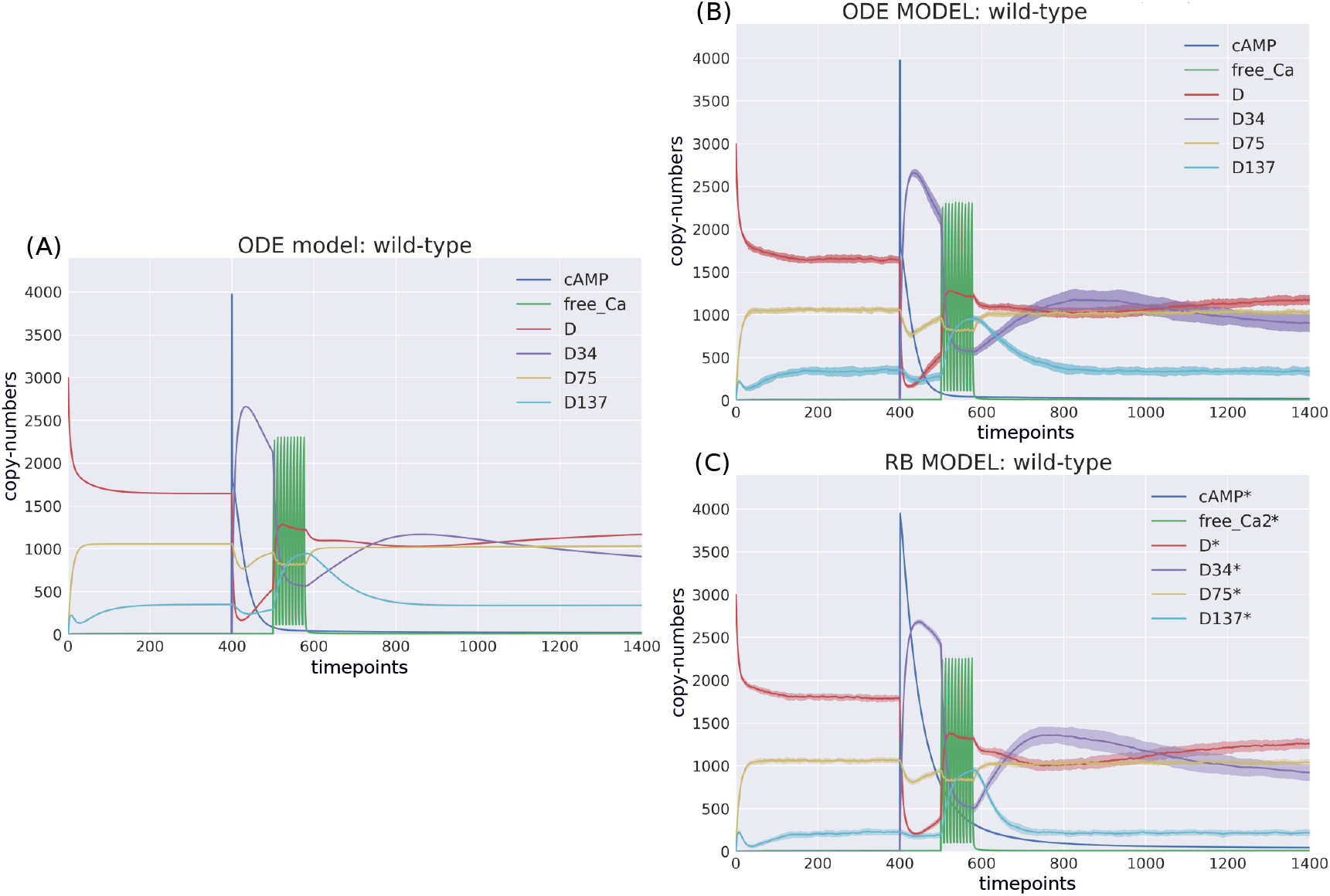
Time-courses of the ODE model for DARPP-32 isoforms triggered by a pulse of cAMP followed by a train of Ca2+ spikes obtained with (A) a deterministic solver, and (B) a stochastic simulation. Trajectories of the stochastic simulation were obtained from calculating mean value (*line*) and standard deviation (*shade*) based on 40 simulations. (C) RB model (stochastic simulation). Variable isoforms of DARPP-32: “D” - unphosphorylated; “D137” - Ser137 phosphorylated; “D75” - Thr75 phosphorylated; “D34” - Thr34 phosphorylated.

For a closer examination, traces of 15 observables (defined in Table 1) obtained from ODE and RB simulations were paired and superimposed (Figure 4). Next to the clear matches (e.g. Figure 4: B, E, H, N), there are discrepancies between paired curves. Five of these 15 observables (Figure 4: C, F, I, J, O) are examples of the largest divergence between models following a similar pattern of behaviour. They are directly connected in a chain of activation reactions that begins with Ca^2+^ (Figure 5). Higher abundance of all Ca^2+^ ions present in the system of the ODE model (Figure 4C) could explain differences between the remaining 4 observables. However, the trajectory of “all_Ca” remains at the 0 level during steady states rising only in the spiking interval, which resembles the abundance of free Ca^2+^ (Figure 4B). Ca^2+^ activates PP2B represented by the trajectory “PP2Bactive”. The higher level of “PP2Bactive” is consistent with the other three observables, suggesting that this is a factor generating the divergences between the models. Based on the curves of the ODE model, we can reason that a stronger activation of PP2B results in proportionally more copies of the unphosphorylated CK1 and phosphorylated D137. This, in turn, increases substrate availability for PP2C; therefore, more copy-numbers of its bound form. This effect is inverted in the trajectories derived from the RB model.

**Figure 4.**
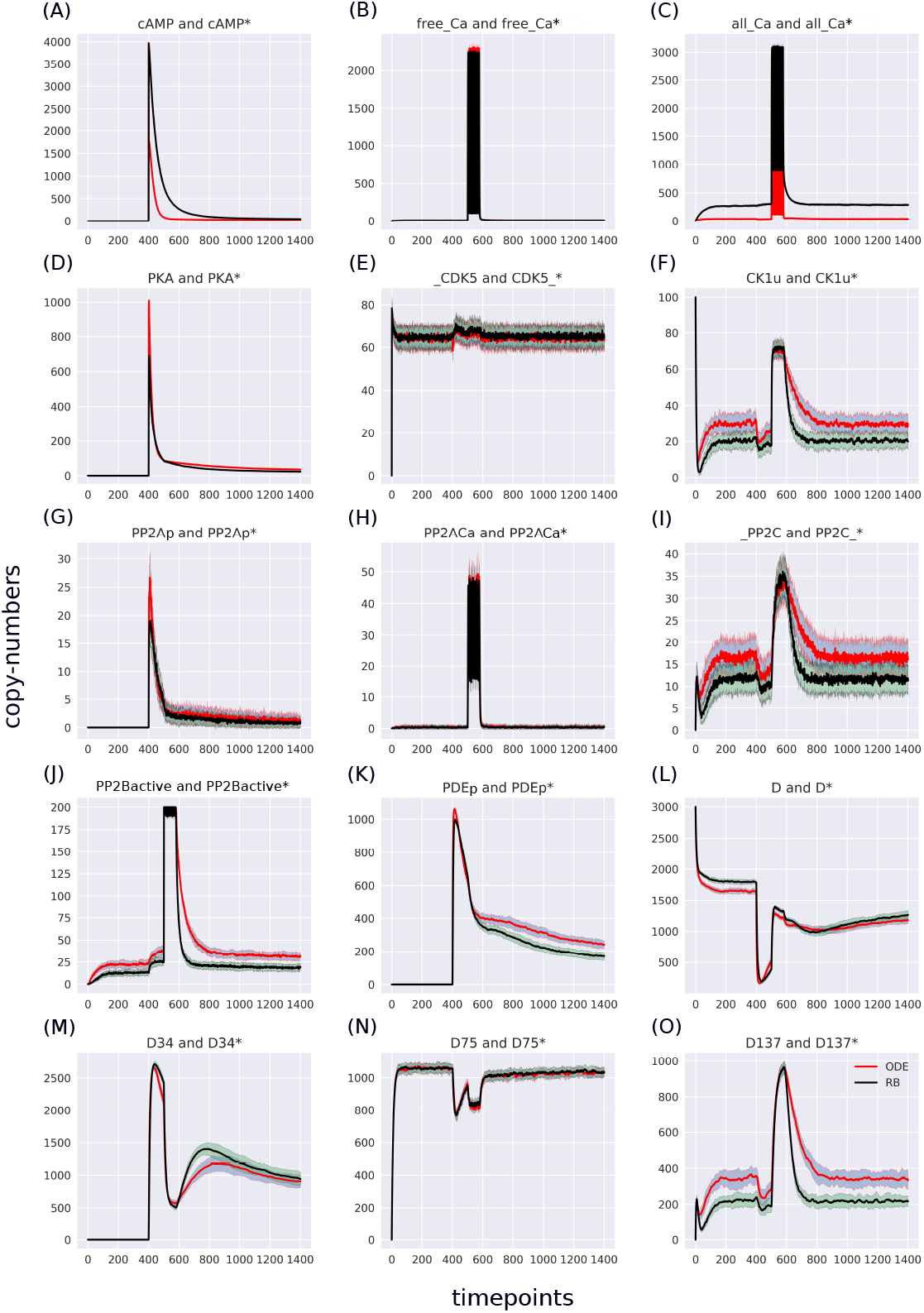
Overlaid time courses of the ODE stochastic and the RB stochastic in the baseline condition. Note that the scales on the y-axis are different to closely compare the traces of the observables. Trace colour: ODE (red), RB (*black*).

**Figure 5.**
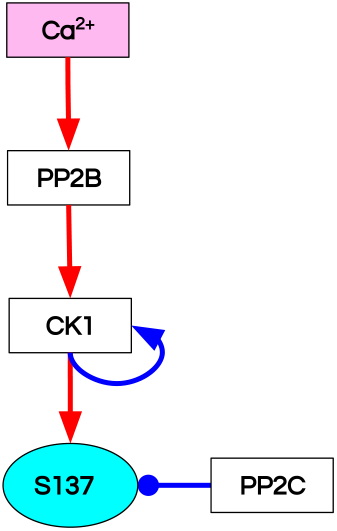
Reaction diagram of the observables showing the greatest divergence between the observables of the ODE and RB models. These observables are connected in a chain of mutually dependent activation reactions triggered by the influx of Ca^2+^.

### RB language allows for detailed dissection of observed molecular species facilitating reuse and reanalysis of dynamic models

It remains unclear why the “all_Ca” observable trajectory produced by the ODE model is much lower than in the RB model at the steady state. Moreover, “PP2Bactive” appears to dictate the higher effect on the other three observables (“D137”, “CK1u”, “_PP2C”). Therefore, “all_Ca” and “PP2Bactive” are closer analysed in further steps.

According to the reaction system underlying both models, the activated PP2B is a complex of 4 Ca^2+^ ions and PP2B. This detail is not explicitly stated in the variable name of the ODE model. Therefore, to obtain the trajectory of all Ca^2+^ ions, the sum of the copy numbers of the molecular species whose variable names contain ”Ca^2+^”, would have to be replaced by a more thorough analysis of the relevant reaction context of the ODE model. This is not the case in the RB model, where an observable of interest is obtained with an automated procedure that sums the trajectories of molecular species containing the specified expression pattern. Since the trajectory of all Ca^2+^ in the RB model includes the ions bound to PP2B, the comparison of “all_Ca” to “all_Ca*” is inaccurate due to a difference in the molecular species included in these observables. A similar inaccuracy, related to the naming of the observables, explains the discrepancy between the time courses of the total number of cAMP observables (Figure 3). In contrast to the RB model, the multiple copies of cAMP bound to R2C2 are not included in this ODE model time course (compare Figure 4A).

As a complete list of molecular species in the RB model is not included in the specification, it is also unknown whether there are other trajectories of molecular species summed up in “all_Ca*”. To obtain these, all molecular species containing Ca^2+^ in the RB model simulation were isolated from snapshot data ammounting to 24 compared to only 13 in the ODE model (“all_Ca”). These 13 species correspond in molecular composition to 18 of the 24 RB species sampled in total. The six absent species in the “all_Ca” observable are composed of an active form of PP2B containing 4 Ca^2+^ ions, either free or bound to phosphorylated CK1, or DARPP-32 in 4 different combinations of phosphorylation states. The number of species comprising “all_Ca” observable in the RB model is higher by five because the half-active form of PP2B (bound to two Ca^2+^ ions) in the RB model exists in 6 variants. Whilst in the ODE model, it is represented as a single species, named “PP2BinactiveCa2”.

Now that we know the composition of molecular species comprising the “all_Ca*” observable feature, we can rerun the RB model with the same components as in “all_Ca” of the ODE model and compare their trajectories directly. To match this newly defined RB observable of “all_Ca*” to the original model, these 18 species were summed to obtain a single observable (Figure 6B). In comparison to the unaltered species composition (Figure 6A), the result shows that discrepancy between the ODE and RB observable trajectories have diminished. Knowing that there is a higher number of forms representing the half-active PP2B in the RB model, we can try to obtain a closer match between the “all_Ca” observables by superimposing only one of 6 trajectories of the RB model. Figure 6C shows that the match is close to perfect. It demonstrates that the differences between the “all_Ca” observables of the two models can be explained by the difference in the number of representations of the molecular species. As the six trajectories have the same dynamics and the same average levels of abundances, choosing one of them is arbitrary (Figure 1). Moreover, the distinction between locations of two Ca^2+^ ions on the numbered sites of PP2B is irrelevant since all 4 sites are functionally indistinguishable.

**Figure 6.**
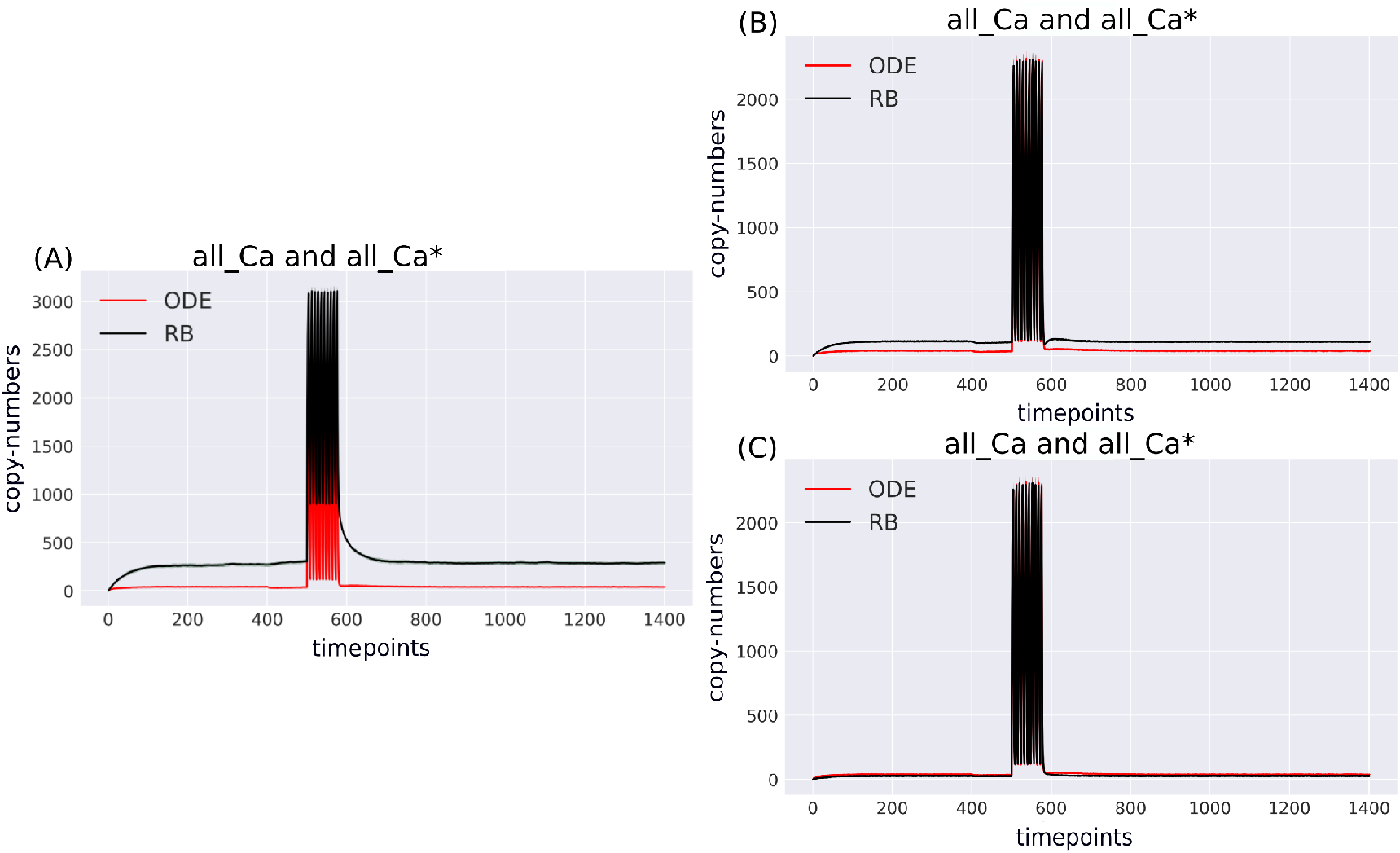
Comparison of variable compositions of molecular species containing Ca^2+^ ions tracked in the system in both models with (B) unaltered observables; (B) all molecular species containing Ca^2+^ ions selected as indicated by names in the original model and summed to obtain a single trace, where 13 molecular species in the ODE model are represented by 18 species in the RB model; (C) 13 molecular species of ODE model matched to 13 of RB model, where only 1 of 6 molecular species of inactive PP2B was selected.

The above analysis demonstrates the flexibility of the RB modelling as a tool to explore complexes formed during the simulation. Molecular species defined in the ODE framework are fixed and definite, whereas, in the RB model, they are a subject of investigation. Snapshots of molecular mixtures permit the determination of created species and their abundances. The results of the RB model can be easily dissected by tracking precisely defined observables of interest during the model run. In addition, the automated agglomeration of time courses into observables renders the RB modelling more advantageous compared to the ODE-based framework. This automation is less error-prone and independent of variable names.

### Rate constants of reactions formulated with “combinatorial-binding” notation should be increased to match ODE trajectories

The largest discrepancy between trajectories can be observed in “PP2BinactiveCa2” (Figure 7M) that for the RB time course was obtained by summing 6 entities representing a half-active PP2B into one. If divided by 6, representing a single variant of half-active PP2B, the trajectory of the RB model becomes lower than the one of the ODE model (Figure 2M). These six forms of the half-active PP2B suggest that a better fit between the two models can be achieved by decreasing the constant rate of rules that represent the binding of Ca^2^ + to free PP2B. However, this could bring the desired effect solely for the half-active PP2B (“PP2BinactiveCa2”) but not for the dynamics of other coupled observables. In particular, the fully active PP2B (“PP2Bactive”), of which the “PP2BinactiveCa2” is an intermediate form. With the current parameter values, the fully active PP2B in the RB model is lower than in the ODE model (Figure 4J). Thus, the decrease in the rate constant of its intermediate form would lead to a further decrease in its copy number. Therefore, although there are more species of the half-active PP2B in the RB model, its fully active form has lower levels than the ODE model. To further examine this observation, we can return to the comparison of model specifications (Table 2). The 4 reactions of PP2B activation are represented by 24 rules, explicit in site-specific detail that includes all combinations of positioning Ca^2+^ on 4 sites of PP2B. Moreover, instead of 3 distinct PP2B species in the reaction representation (inactive, semi-active, activated), in the rule-based model there are 8 different molecular species. Thus, the fine-grained representation of species in the molecular mixture slows down the transition from inactive to active PP2B, despite the two-step transition encoded in both formalisms. The same can be observed in the second example that required a much larger number of rules, i.e. an activation of PKA. Figure 4D shows that the RB trajectory of the “PKA*” observable also reaches a much lower peak than its ODE counterpart. Accordingly, values of rate constants of rules that the number increased due to the “combinatorial binding” notation in the RB model, i.e. are represented by more variants of species, should be increased to closely match the ones in the ODE model.

**Figure 7.**
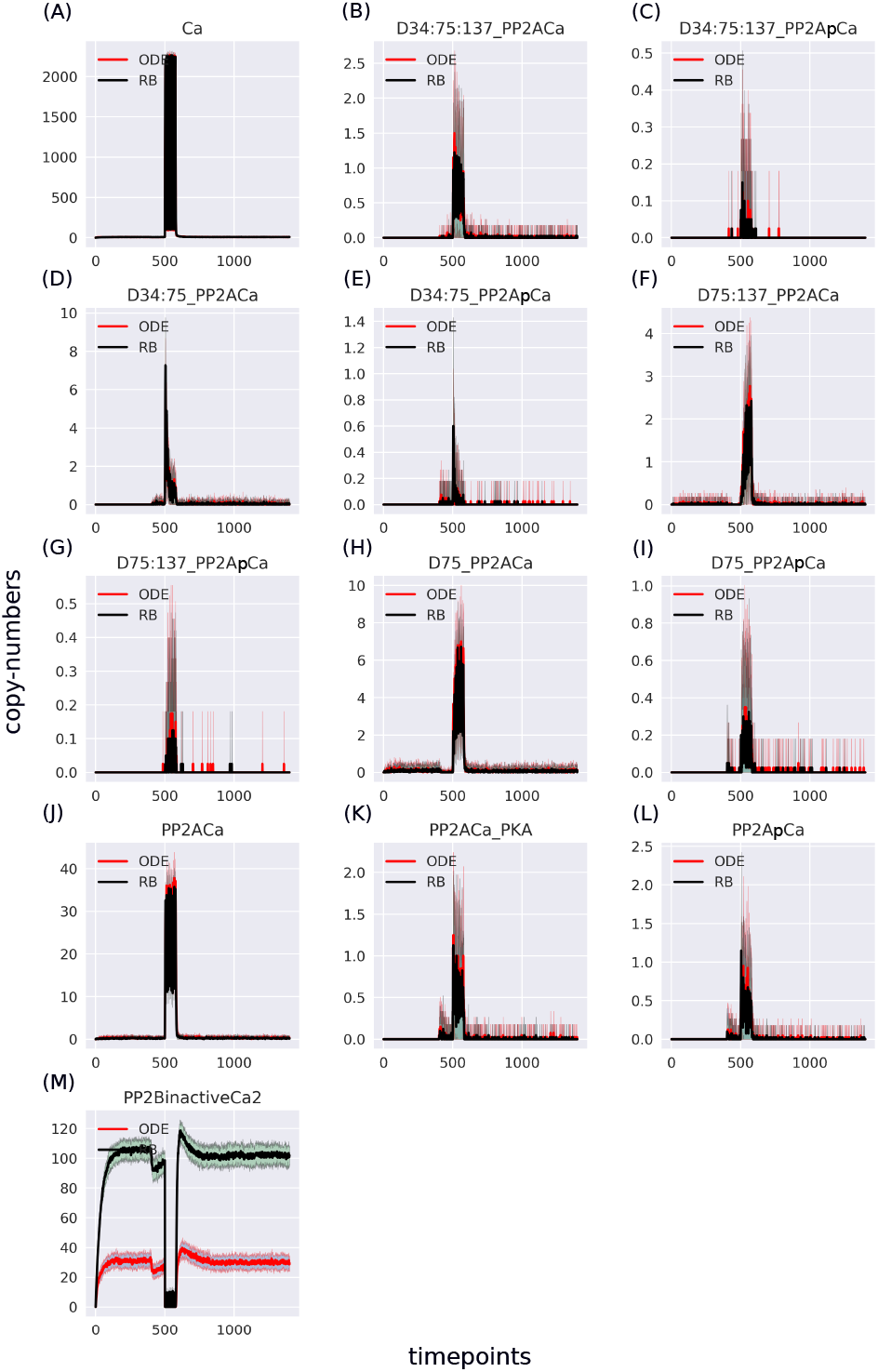
Traces of 13 pairs of molecular species containing Ca^2+^, selected to match the ODE model. The largest disparity lies in the “PP2BinactiveCa2” variable - summation result of 6 entities representing an inactive form of PP2B in the RB model.

### RB language facilitates modifications of dynamic models

#### Site-directed mutations

The authors of the ODE model analysed it in two other conditions of site-directed mutagenesis affecting the Ser137 site. The same perturbations were applied to the RB model to establish if they would generate similar results. Figures 8 juxtapose simulation results of both models affected with constSer137 and Ser137Ala mutations. As exemplified by the six key observables, there is a close match in initial conditions and a general pattern of dynamics between the time courses of the two models. Therefore, we can conclude that the RB modelling allows emulating experimentally observed perturbations similarly to the ODE-based modelling.

**Figure 8.**
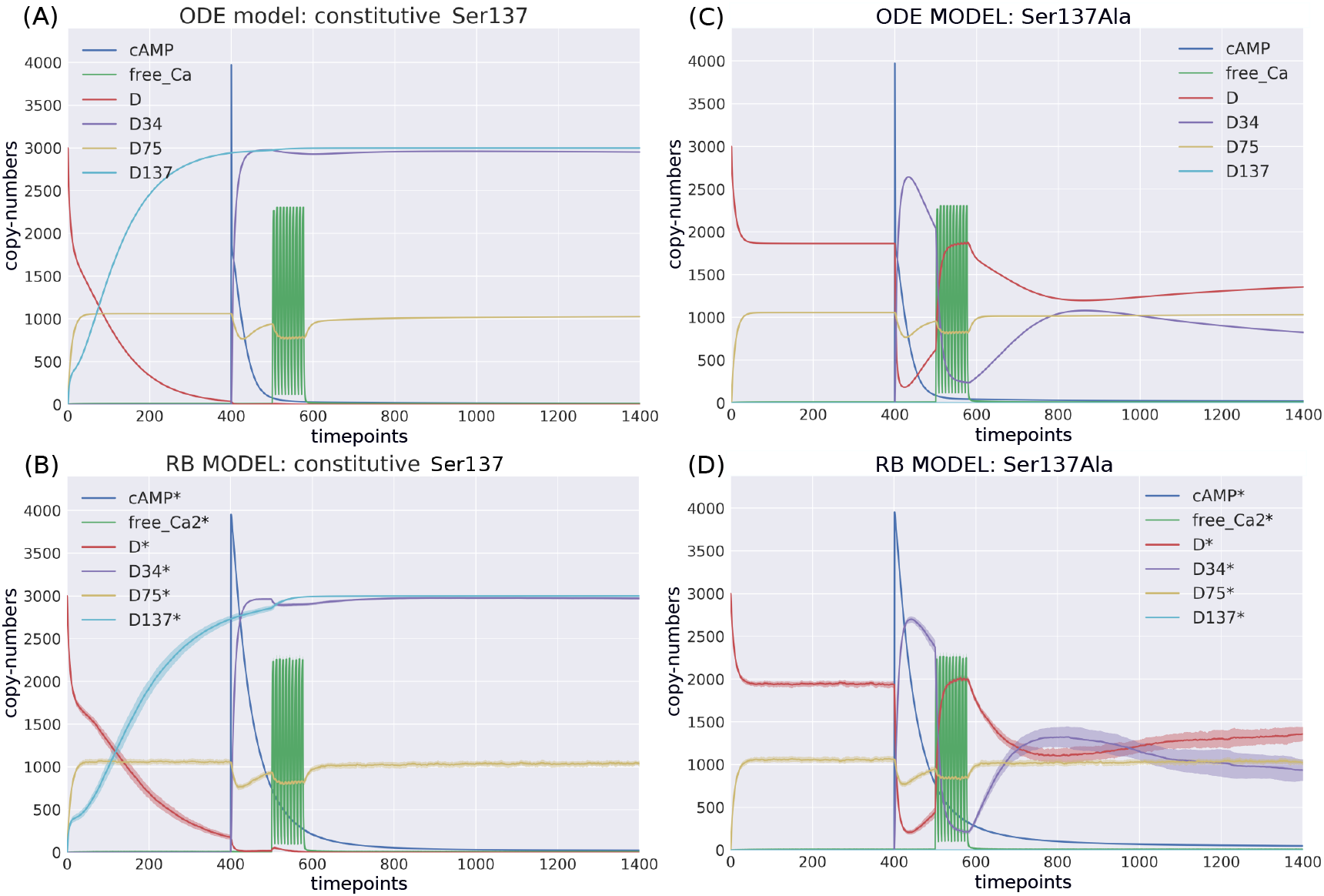
Comparison of the constitutive Ser137 mutation induced in (A) ODE model in deterministic setting; (B) RB model in stochastic setting; and the Ser137Ala mutation in (C) ODE model in deterministic setting; (D) RB model in stochastic setting; The same interference performed on rate constants of the two models caused similar dynamics.

#### Competitive and non-competitive site binding

Two variants of the RB model with different binding site specifications are compared to test whether the dynamics of the model are affected when DARPP-32s binds multiple partners at once. The first specification is a competitive variant of the model with one binding site (oBS). In the second, a non-competitive variant, the partners bind simultaneously (three-binding-sites, tBS). This type of comparison between ODEs and RBs was previously presented by Blinov et al. (2006). In contrast to their results, the superimposed trajectories of two models (Figure 3) demonstrate no effect on model dynamics. As the direct consequence of this modification is an increase in the size of the complexes to more than two proteins, it seems that larger complexes are rarely formed during the simulation. This interpretation is confirmed in a direct examination of species counts (Figure 4). As these three binding sites do not counter each other’s binding properties, the lack of difference might be caused by the similarity in occupancy between a single site and all three sites together. The probability of a site being connected depends on copy numbers of reactants and the strength of binding affinities. Reactions in the model are classified as weak, with dissociation rates in the range of *μ*M. Low-affinity bindings generally lead to lower levels of site occupancy. Moreover, the amount of DARPP-32 molecules exceeds the total counts of all its interactors. Therefore, with the current proportions of reactants, all three sites of DARPP-32 cannot be saturated to expose the difference in the binding capacity of DARPP-32. To expose the potential differences in dynamics between two binding scenarios, the site occupancy must also be modified. The simplest test of this explanation could be performed by altering the size of the reactant pools. For instance, a significant decrease in DARPP-32 levels could increase the proportion of other reactants.

## DISCUSSION

ODE-based modelling is a classical and commonly used method for creating detailed dynamic models of biological systems (Lotka, 1920; Hodgkin and Huxley, 1952; Lisman, 1989). It is frequently a point of reference and comparison to newly proposed modelling methods (Morris et al., 2010; Ciocchetta and Hillston, 2008; Chaouiya, 2007; Chylek et al., 2015; Danos and Laneve, 2004). Nevertheless, modelling of signalling systems with ODE poses difficulties due to complexities underlying molecular interactions (Kholodenko, 2006; Stefan et al., 2014). RBM was proposed as a solution to this problem.

Numerous reviews (Danos, 2007; Chylek et al., 2014; Hlavacek et al., 2006) and studies discussed the advantages of RB modelling over ODE. This paper presents results of encoding reactions underlying an ODE model to the RB language and a comparison of their specification and simulation results. The manual translation of this ODE model into any RB language was necessary despite the existence of a method (Tapia and Faeder, 2013) for automated translation of the SBML-format encoding ODE-based models to an RB model format. The translation failed to generate executable model with correctly identified agents (Supplement 1).

### Effects of the RB framework on the model notation

Encoding reactions into rules slightly reduced the size of model specification and increased counts of molecular species, which confirmed the well-known advantage of rule representation (Hlavacek et al., 2006; Chylek et al., 2014). Closer analysis of reaction subsets representing more general molecular mechanisms showed that reduction in the reaction number is only true for reactions occurring between the same reactants, describe the same transformation, and are parameterised with the same values of rate constants, but differ only concerning binding or internal state of reactants. In this type of reaction, the number of unique reaction rates was equal to the number of rules. An increase in the number of rules representing reactions occurred where the same partner binds an agent at multiple sites. In such reactions, all possible positions and stages of the binding process had to be explicitly encoded. The “combinatorial binding” notation is not a general property of the RB language but characteristic of the Kappa syntax. In the alternative to Kappa RB framework, BNG, a rule is definable with identical sites’ names. This implies that the rule pattern defined for one applies to the others, effectively shortening the rule description (Sekar and Faeder, 2012).

### Discrepancies in dynamics between models

As reported in other similar study (Blinov et al., 2006), comparison of trajectories showed an agreement between models’ dynamics, with some discrepancies. It should be noted that though there is an issue of ambiguous molecularity in Kappa, as opposed to BNG (Feret and Krivine, 2012), this problem does not arise in the DARPP-32 model, as none of the rules allow for closed rings between agents. These discrepancies mainly appeared due to a lack of precision in the variable names of the ODE model caused by differences in molecular species comprising tracked observables. When the simulation of the RB model was performed with observables exactly matching the ones in the ODE model, almost all paired trajectories fitted perfectly. The only problematic observable involved the “combinatorial binding” notation. Further analysis suggested that activation of proteins encoded with fine detail is slower than in the ODE model. It is worth noting that although the use of BNG removes the need to define all possible bond combinations explicitely, this does not mean that it can change the results of Kappa simulations This claim can be supported with an example given by the developers of BNGL in a publication reporting on the translator between BNG and Kappa, TRuML (Suderman and Hlavacek, 2017). Their example shows on antibodies with identical antigen-binding sites that equivalent rule sets in both RBM frameworks have the same rate constants. In the BNG framework, however, this rate constant is automatically scaled by a factor equivalent to the multiplicity of indistinguishable ways to obtain the reaction product (Faeder et al., 2005, 2009). The authors’ comparison of the time traces of Kappa- and BNGL-defined rules shows equivalent results under stochastic realisations (Suderman and Hlavacek, 2017). An earlier study by Suderman and Deeds (2013) also showed equality of results between both frameworks.

Thus, it can be argued that the results of such a comparison carried out for our model will be the same, because the molecular mixture when simulated with NFsim (Sneddon et al., 2011), a variant of the network-free simulator for BNGL, will implicitly contain the same number of molecular species with different binding variants as it is in KaSim. Thus, we argue that “combinatorial binding” expressed as a single rule in BNGL will not change the dynamics observed with Kappa. From a formal modelling perspective, this effect is understandable because the fine representation of the agents’ states expands their probabilistic distribution, increasing the probability of staying in an intermediate state much longer. These specific discrepancies should be accounted for when reactions and rate constants from ODE models are reused to build RB models. Nevertheless, the experimental rates for these types of reactions need to be determined and compared.

### A shift in modelling focus with the use of the RB framework

The process of encoding reactions into rules turns attention to questions such as how many binding partners can simultaneously bind a protein. The translation process has shown that information about interfaces of interacting proteins and their alternative states would considerably ease the process of model development by guiding decisions on agents’ signatures. Therefore, data resources that could support RB modelling, such as a source of protein interaction interfaces, post-translational modifications (PTMs), and protein domains. For instance, proteins containing phosphatase catalytic domains are enzymes of dephosphorylation reactions (Sacco et al., 2012). However, such detailed information is not accessible for most molecular agents.

### Facilitation of modification and reuse of dynamic models

Snapshots, in other words state vectors sampled during the simulation, have been used as a tool for model exploration in other studies (Sorokina et al., 2011; Suderman and Deeds, 2013). Snapshot visualisation is included as a standard tool in the Kappa framework (Boutillier et al., 2018). We used snapshots to demonstrate detailed exploration of the emerging molecular species during the simulation of an RB model. By overlaying trajectories of molecular species we could track the source of differences between models. Emerging molecular species are relatively easy to examine and analyse. Unlike Kappa, in BNG it is possible to count the number of possible species through network generation. However, this option reaches its limit when the model has a significant size and network generation is not possible. Then, if it is not sampled by simulations, it is calculated analytically (Suderman and Deeds, 2013). While being an approximate approach to species counting, snapshots provide sufficient estimates and measures to examine the system.

Conversely ODE models rely heavily on complete knowledge of the system and the arbitrary naming of variables. Retrieval of molecule counts, hidden in individual molecular species of the ODE model, would require further deconstruction of the reaction system and the arduous extension to much more complex models. The implicit difficulty of identifying all molecular entities arising from the molecular species in ODE models fundamentally impacts on their veracity.

The precise identification of molecules among species could be performed by parsing the SBML-based model encoding. However, exploration of the Fernandez et al. (2006) model web page in the BioModels website^1^, shows incomplete annotations of molecular species behind variable names, both concerning the actual counts of interactions (e.g. Ca^2+^) and their components. In this light, automated identification of molecules in the RB framework is particularly advantageous, as it offers a transparent framework with error-prone identification of molecules in a modelled system. This feature is particularly vital when the modelled system is composed of numerous molecules, and their particular states are to be analysed in detail.

The RB model was tested with two types of site-directed modifications, demonstrating the framework’s flexibility to reproduce experimentally conducted perturbations. Though the binding site modification effectively changed the model reaction network, it did not affect the model response. Nevertheless, this intervention demonstrated the ease of performing such alterations within the RB framework. Additionally, the pattern notation improves model clarity and provides an intuitive representation of a model akin to a set of chemical reactions rather than equations, potentially improving the learning curve for a modeller-to-be.

### Limiting factor of the simulation time

The execution time of stochastic model simulations in comparison to deterministic ones has always been an issue addressed by multiple optimisation strategies (Gillespie, 2007). It is no different in the case of the two models compared here. It takes almost 40min^2^ to simulate this particular RB model with the KaSim simulator. The solution of the ODE model in the COPASI environment in the deterministic setting returns in an instance. The COPASI stochastic simulator produces results in no more than 15sec. Thus, reducing simulation times for RB models remains an important challenge for use cases in which speed is critical. However, with a sufficiently simple model, network-based simulation can be performed far more efficiently using BNGL, keeping in mind that more complex models will have a higher runtime with network-based simulation (Sneddon et al., 2011). When generating large-scale models, the type of RB model simulation can be scheduled on a case-by-case basis (Santibáñez et al., 2020).

### Further explorations of the DARPP-32 RB model

There are two main routes for further exploration of the RB model. The first one is a modification of parameters defining different phases of combinatorially bound Ca^2+^ ions to PP2B and cAMP to R2C2. A particular task would be to identify factors by which the binding constants could be modified to counteract the many intermediate variants of these complexes and the lower copies of their activated final forms. It would be interesting to identify conditions under which we could observe a difference in model dynamics after adding binding sites to DARPP-32. As a starting point, the modification could be achieved by significantly decreasing the copy number of DARPP-32 compared to other interactors. As mentioned by Fernandez et al. (2006), levels of DARPP-32 vary considerably, between *μM* to tens of *μMs*, in the striatum. With the greater availability of single-cell techniques for protein quantification (Lo et al., 2015) it would be worth establishing more precisely the range of DARPP-32 even at the resolution of a dendritic spine (Otmakhov and Lisman, 2012). Estimating variability between cells could also be used to compare the varying levels of phosphorylated DARPP-32 at Thr34 that were observed in the stochastic simulations.

### Extending the model to match evolving knowledge on the DARPP-32 interaction network

The main advantage of RBM over ODE is the flexibility to extend and manipulate the model, rather than rewriting it into a new model with other limitations due to the type of expressivity of ODEs. In this way, the community interested in molecular systems related to DARPP-32 can pool their knowledge in a single representation. We believe that the refactored model of the DARPP-32 in a rule-based framework will facilitate the creation of such an updated model in future. It should be noted that the ”DARPP-32 events” pathway (R-HSA-180024.3) specified in the REACTOME database is in fact the model of Fernandez et al. (2006) used in this study. So far, only early signalling events of DARPP-32 have been modelled, localised mainly in the cytosol. Thus, the Ser102 site was omitted in our model because there is no evidence that it can be affected by DA or Glu signalling (Girault et al., 1989; Svenningsson et al., 2004). However, a recent study by Nishi et al. (2017) suggests that Glu can decrease the effect of DA signalling (phosphorylation of three other sites) by dephosphorylating DARPP-32 at Ser102 causing the accumulation of DARPP-32 in the nucleus. The Ser102 site regulates nuclear transportation of DARPP-32 (Stipanovich et al., 2008) representing late signalling events. Stipanovich et al. (2008) have shown that nuclear accumulation of DARPP-32 is promoted by drugs of abuse; the inclusion of Ser102 in future models could be valuable to further explore this. Additionally, to calibrate the dynamics of the 4 phosphorylation sites, one could use a recently developed tool for RB models, Pleione (Santibáñez et al., 2019), taking into account measurements obtained by Nishi et al. (2017).

The inclusion of more than two major phosphorylation sites in modern DARPP-32 models can be further justified by the fact that the other three sites (excluding Th75) may influence the phosphorylation of Thr34, the most important functional site (Girault and Nairn, 2021). Furthermore, the role of the DARPP-32 interaction network in the nucleus appears to be an important modelling target in a recent model incorporating DARPP-32 (Yapo et al., 2018), indicating the need to clearly demonstrate the contribution of DARPP-32 to the observed switch-like action of PKA in the nucleus. This model was written as an ODE and takes into account the action of two of the four DARPP-32 phosphorylation sites. However, since the Ser102 site regulates nuclear transport of DARPP-32, it will have to be included in such models. The resulting complexity of the model may be difficult to capture as an ODE.

Although it was not our aim to reveal whether stochastic simulation can demonstrate new behaviour of this particular molecular system that deterministic simulation has so far not revealed, this would be also valuable to address in future research.

### Need for formal prioritisation methods of emerging molecular species

RB modelling offers tools for the dissection of emerging molecular species during simulations. As the number of such molecular species increases it becomes increasingly difficult to identify which model components are of particular importance to the system. There is great potential to exploit commonly used methods for model exploration, such as sensitivity analysis, to identify critical model features including parameters and model output variables. It would be advantageous to support the modeller’s assumptions with automated methods to prioritise model outputs for downstream analysis and greater insight into the underlying biological systems.

## CONCLUSIONS

Dynamic molecular modelling has become increasingly important in uncovering and integrating our dispersed knowledge of molecular mechanisms. Choosing the best formalism meet this challenge is a difficult task. In this work, we have presented a detailed and systematic comparison of two major formal approaches to quantitative modelling. We demonstrated the advantages and disadvantages of RB modelling to the preeminent ODE approach. We confirm, after other similar studies, that RB modelling is more detailed and flexible way to represent biological molecular systems, enabling exploration of individual molecular entities, model extension, and future reuse. These conclusions confirm the huge potential of the RB formalism and hopefully will embolden future exploration and research in this topic.

## CODE ACCESSIBILITY

The code reproducing the figures and the encoding of the Kappa models can be found on github : https://github.com/ewysocka/rb_vs_ode_model_of_darpp-32.

## LIST OF ACRONYMS

BNG: BioNetGen
Ca^2+^: calcium ions
cAMP: cyclic adenosine monophosphate
DA: dopamine
DARPP-32: dopamine- and cAMP-regulated neuronal phosphoprotein with molecular weight 32 kDa
EGFR: epidermal growth factor receptor
Glu: glutamate
MSPN: medium spiny projection neurons
ODE: ordinary differential equation
PTM: post-translational modification
RB: rule-based
RBM: rule-based modelling
SBML: Systems Biology Markup Language
Ser102: Serine 102
Ser137: Serine 137
SSA: Stochastic Simulation Algorithm
Thr34: Threonine 34
Thr75: Threonine 75

## SUPPLEMENTARY MATERIALS

### 1 AUTOMATED TRANSLATION OF ODE MODEL WITH ATOMIZER

BioNetGen (BNG) is a member of the family of rule-based formalisms that most closely resemble Kappa. The BNG framework is supported by the automated translation of SBML into BNGL format by Atomizer (Tapia and Faeder, 2013). Atomizer defines molecular bond structures, implicit in reaction-based models, through the use of molecular species naming conventions and reaction stoichiometry. This method could potentially offer an easy way to derive an RB model from the original ODE. The performance of the Atomizer has been tested on the Fernandez et al. (2006) model to compare the results of this automatic translation with a manual translation of the model. The automatic translation was successful, but simulation of the resulting model resulted in errors, regardless of using all available model simulation options. The standard error output reported many instances of conflicting definitions and inconsistent naming. In notational terms, the complexity and redundancy of the model coding appeared not to be designed to be edited by a human and therefore it was difficult to assess correctness without simulating the model. The generated model was examined in terms of agent and rule definitions. Examination of the rules revealed fully contextualised reaction instances (Code 3). All combinations of DARPP-32 states were represented as separate species (Code 1, l.2 and l.3).

These results supported the need for a manual translation of the ODE model into Kappa.

**Code 1.**
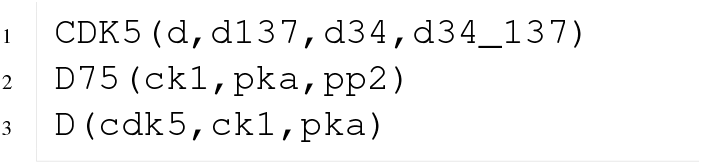
Example of agents in BNGL generated with Atomizer

**Code 2.**
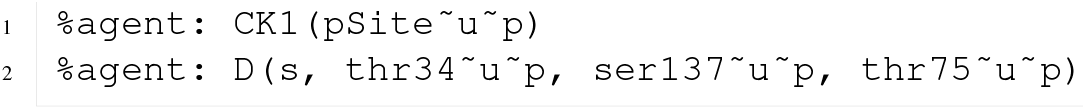
Example of agent manually formulated in Kappa

**Code 3.**
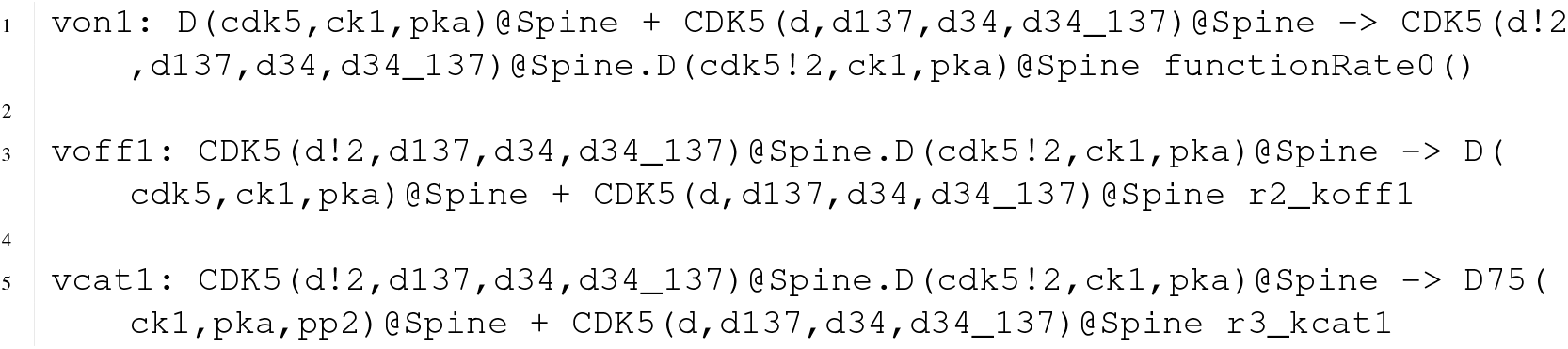
Example of rules in BNGL for a two-step phosphorylation generated with Atomizer

**Code 4.**
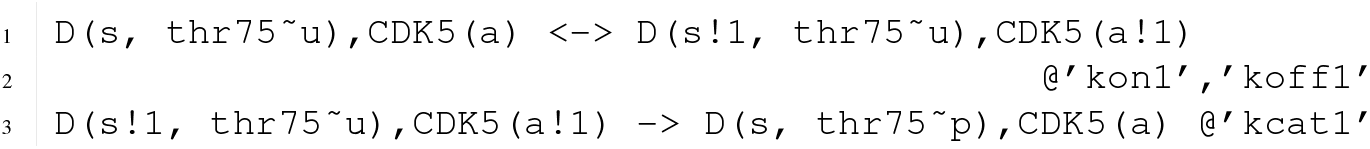
Example of rules for a two-step phosphorylation manually encoded in Kappa

**Table 1.** Names of RB observables and corresponding names of ODE observables with definitions. To obtain observable ODEs, the time series of the corresponding molecular species are summed based on their names.

| RB | ODE | Definition |
| --- | --- | --- |
| cAMP* | cAMP | cAMP binding unspecified |
| free_Ca* | free_Ca | Ca<sup>2+</sup> unbound |
| all_Ca* | all_Ca | Ca<sup>2+</sup> binding unspecified |
| PKA* | PKA | PKA binding unspecified |
| CDK5_* | _CDK5 | CDK5 bound |
| CK1u* | CK1u | CK1 unphosphorylated, binding unspecified |
| PP2Ap* | PP2Ap | PP2A phosphorylated, all bindings unspecified |
| PP2ACa* | PP2ACa | PP2A bound to Ca<sup>2+</sup>, phosphorylation and other bindings unspecified |
| PP2C_* | _PP2C | PP2C bound |
| PP2Bactive* | PP2Bactive | PP2B active, binding unspecified |
| PDEp* | PDEp | PDE phosphorylated, binding unspecified |
| D* | D | DARPP-32 unphosphorylated at all sites, binding unspecified |
| D34* | D34 | DARPP-32 phosphophorylated at Thr34 with unspecified binding, other sites' internal states and binding unspecified |
| D75* | D75 | DARPP-32 phosphophorylated at Thr75 with unspecified binding, other sites' internal states and binding unspecified |
| D137* | D137 | DARPP-32 phosphophorylated at Ser137 with unspecified binding, other sites' internal states and binding unspecified |

**Figure 1.**
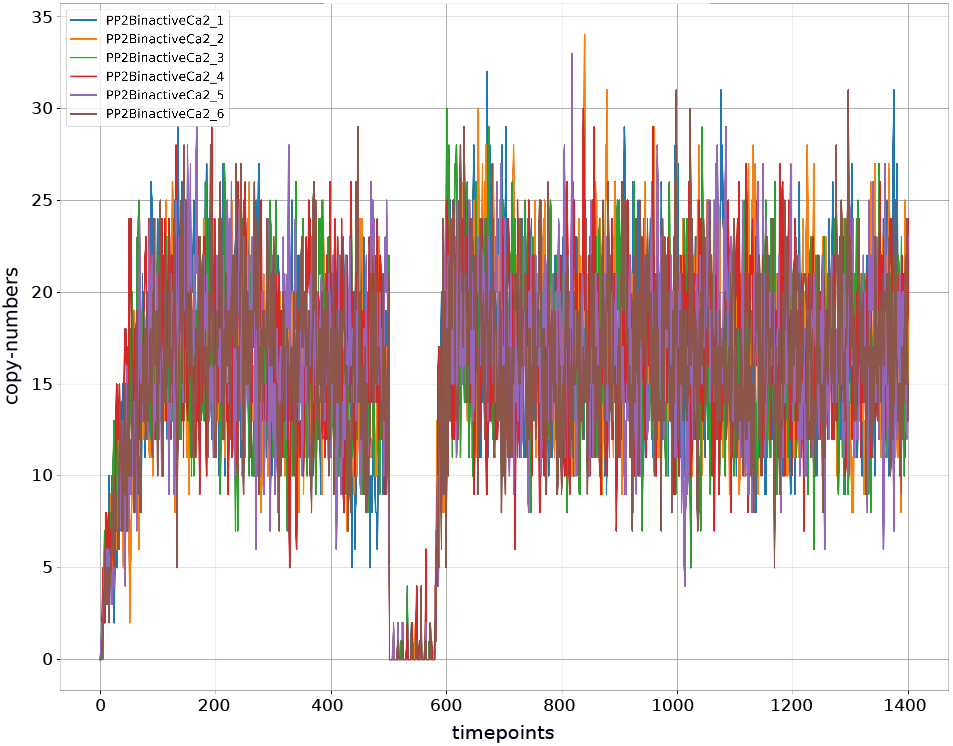
Semi-active PP2B is a complex composed of PP2B and two Ca^2+^ ions. Simulation of the RB model generates six different molecular species representing this complex due to combinatorial binding of Ca^2+^ ions to four identical PP2B sites. The graph shows the superimposed trajectories of these six variants of semi-active PP2B. None of these six trajectories is distinguished from the others by either the pattern of dynamics or the average abundance level.

**Figure 2.**
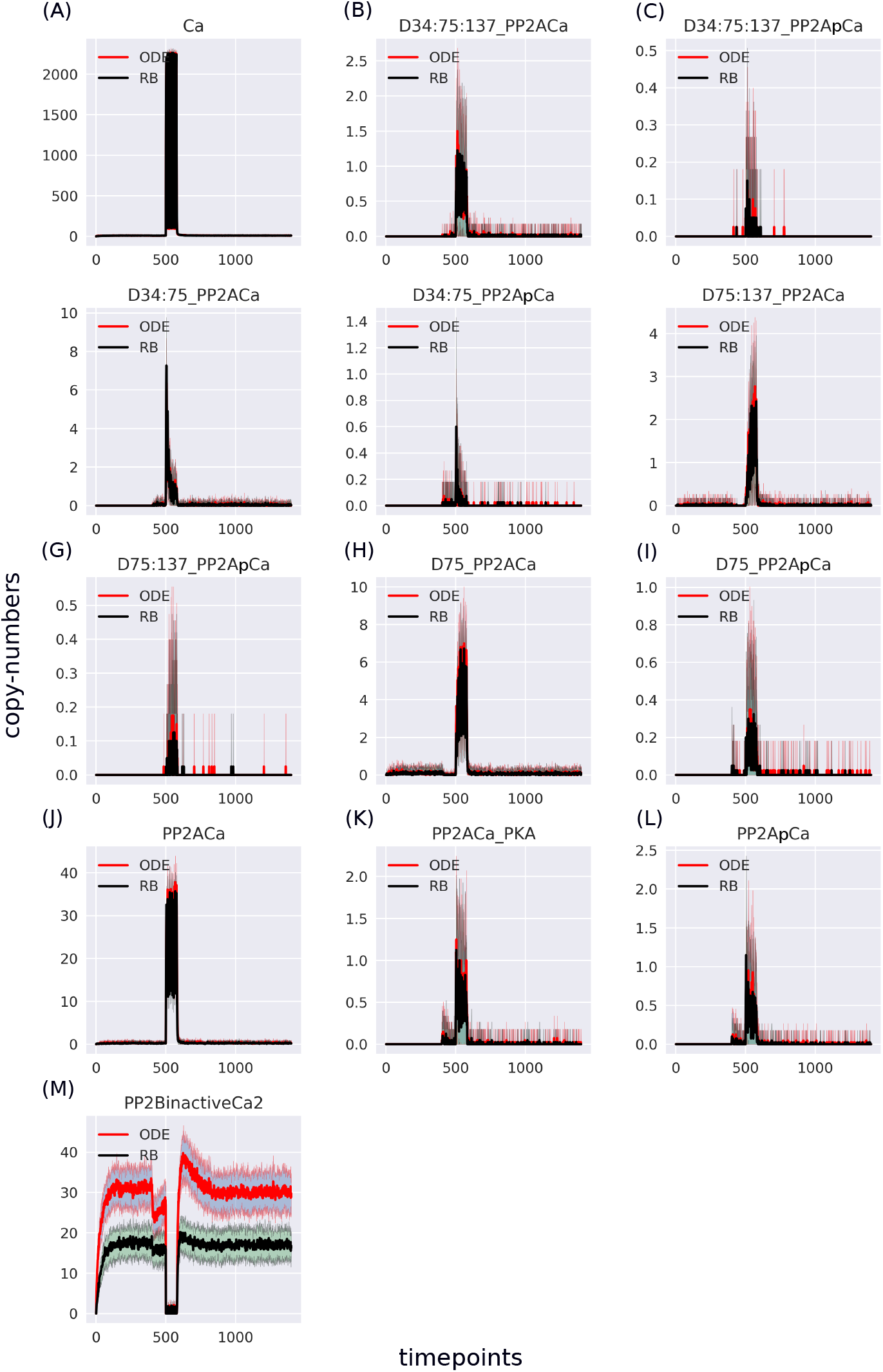
Comparison of separated Ca^2+^-containing molecular species selected as in the ODE model. The “PP2BinactiveCa2” trajectory in the RB model was obtained by selecting one of 6 entities representing, among others, the inactive form of PP2B in the RB model. There is still a discrepancy between the models, but the trajectory is lower for the RB model.

**Figure 3.**
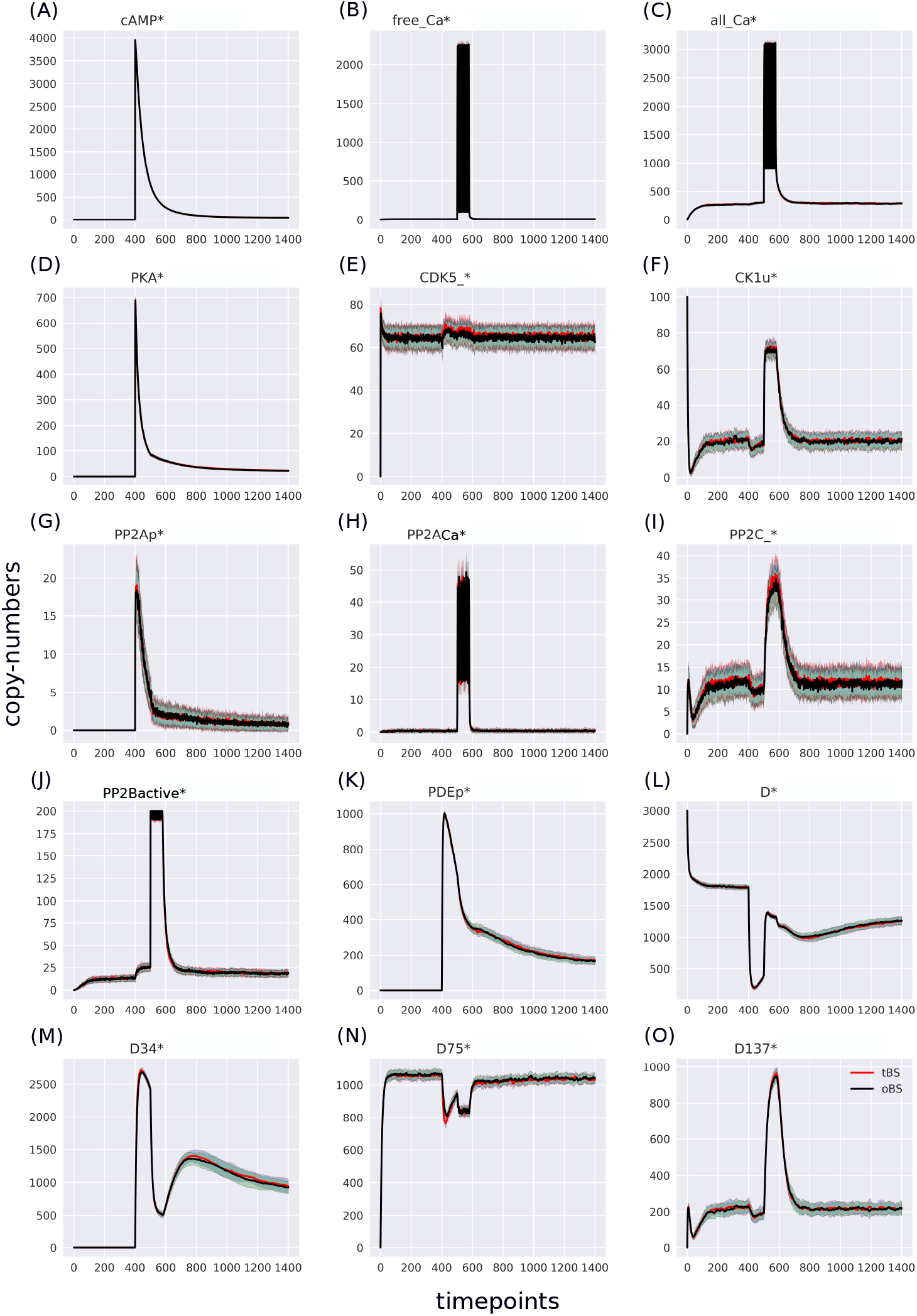
Comparison of two variants of the RB model in which the agent representing DARPP-32 had one binding site (oBS, red trace) and three binding sites (tBS, black trace). The superimposed trajectories of the respective agents indicate that the model trajectories were not affected by this modification.

**Figure 4.**
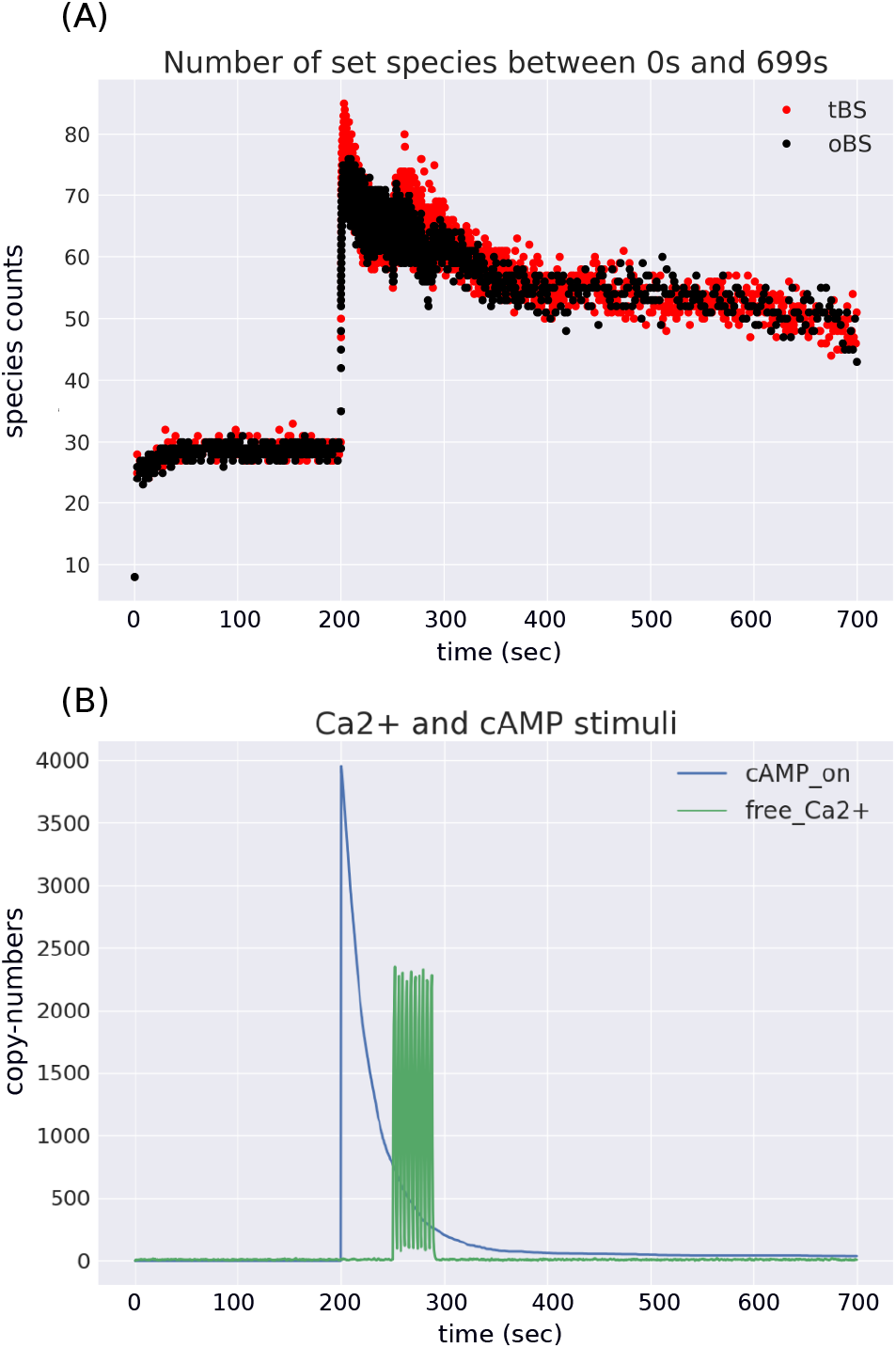
Overlay of changes in the number of species over time for two models in which DARPP-32 can bind at one site (oBS) and three sites (tBS) (A). The size of the species set is similar in both model variants. The change in the number of unique species is consistent with the stimulus trajectory (B). The dynamics of complex formation are dictated by the pattern of stimulus input, as the largest differences between oBS and tBS occur during stimulus application.

## Footnotes

1 http://www.ebi.ac.uk/biomodels-main/BIOMD0000000153

2 The total CPU time measured with the “`time`” command on Linux OS

